# Single-cell RNA-sequencing reveals thoracolumbar vertebra heterogeneity and rib-genesis in pigs

**DOI:** 10.1101/2021.05.25.445704

**Authors:** Jianbo Li, Ligang Wang, Dawei Yu, Junfeng Hao, Longchao Zhang, Adeniyi C. Adeola, Bingyu Mao, Yun Gao, Shifang Wu, Chunling Zhu, Yongqing Zhang, Jilong Ren, Changgai Mu, David M. Irwin, Lixian Wang, Tang Hai, Haibing Xie, Yaping Zhang

## Abstract

Thoracolumbar vertebra (TLV) and rib primordium (RP) development is a common evolutionary feature across vertebrates although whole-organism analysis of TLV and RP gene expression dynamics has been lacking. Here we investigated the single-cell transcriptomic landscape of thoracic vertebra (TV), lumbar vertebra (LV), and RP cells from a pig embryo at 27 days post-fertilization (dpf) and identified six cell types with distinct gene-expression signatures. In-depth dissection of the gene-expression dynamics and RNA velocity revealed a coupled process of osteogenesis and angiogenesis during TLV and rib development. Further analysis of cell-type-specific and strand-specific expression uncovered the extremely high levels of *HOXA10* 3’-UTR sequence specific to osteoblast of LV cells, which may function as anti-*HOXA10*-antisense by counteracting the *HOXA10*-antisense effect to determine TLV transition. Thus, this work provides a valuable resource for understanding embryonic osteogenesis and angiogenesis underlying vertebrate TLV and RP development at the cell-type-specific resolution, which serves as a comprehensive view on the transcriptional profile of animal embryo development.

## Introduction

In vertebrates, vertebrae develop and segment during early embryogenesis [1]. During this process, cervical (CV), thoracic (TV), lumbar (LV), sacral (SV), and caudal (CAV) vertebra are formed in sequence along the anterior-posterior axis [2,3]. Body region allocation and the transition between regions have important morphological, physiological, and evolutionary consequences, given that their relative proportions vary widely among vertebrates [4]. Partitioning of the body into TV and LV has been of long-term biological research interest and many pioneering studies have attempted to identify genes and genomic variations underlying this developmental process. For examples, *OCT4* and *GDF11* are identified as important genes and their overexpression or knockout lead to TV elongation, similar to the long TV partition seen in snakes [5,6]. At the same time, the identity of the thoracolumbar vertebra (TLV) transition is shaped by members of the *HOX* gene cluster in mice [7–9]. Therefore, despite the valuable insights into TLV transition and rib-genesis at the single gene level, previous transcriptomic analyses have not resolved the tempo-spatial gene expression behind these developmental processes. Thus, profiling of gene expression in TV and LV body partitions should offer an opportunity to gain deeper insights into these developmental processes.

The development of single-cell RNA sequencing (scRNA-seq) technologies has provided an opportunity to investigate tempo-spatial gene expression during embryo development. scRNA-seq methods have higher gene expression resolution than traditional transcriptome analysis such as whole embryo transcriptomic sequencing and bulk-segregant sequencing (Bulk-seq) [10,11]. scRNA-seq has an advantage in detecting cell types and the gene expression signatures for each type [12,13]. For instance, cell atlases for mammalian systems including neonatal rib and bone marrow stroma have been generated by analyzing both fetal and adult mouse tissues [14–16]. Cell atlases characterization and gene expression pattern analysis should provide valuable information on the difference during TV and LV development.

Previous studies on TV and LV development have largely focused on mouse models by examining the effects of genetic variation on phenotypes through the over-expression or knockout of genes [7–9]. These models have greatly advanced our knowledge on TV and LV partition development. Alternative models for TV and LV development studies include some domestic animals with varying numbers of TV and CV, such as pigs and sheep [17,18]. These domestic animals may function as valuable model species for further exploration of genes and signaling pathways underlying TLV development with a low genomic divergence among individuals within a species.

In this study, we used the pigs as a model and attempted to explore the cell compositions in the developing TLV and RP. We conducted a single-cell transcriptiomic analysis of cells collected from TLV and RP from one Large White (LW) pig embryo at 27 days post-fertilization (dpf), which corresponds to the commencement of rib formation. Overall, this study provides a rich resource that can advance our understanding of TLV transition and RP development in vertebrates.

## Results

### Identification of cell types and states in developing TV and LV in pig embryo

To gain an insight into the development of TV and LV, we started an analysis by characterizing cell populations from two different anatomical body partitions. To determine the time point for cell sampling, we examined the development of pig embryos at 15, 16, 17, 18, 20, 25, 27, and 29 days post-fertilization (dpf). Our analysis revealed that ribs commenced stemming from TV at 27 dpf, while embryos less than 27 dpf did not show evident ribcages. Ribcage development was completed at 29 dpf [Li et al. unpublished data], therefore, embryo at 27 dpf was used for cell population characterization in developing TV and LV.

A total of 360 (180 TV cells and 180 LV cells) cells from three vertebras close to the TLV segmentation joint were isolated by micromanipulation and enzyme digestion from LW embryo at 27 dpf (**Figure 1**A and Figure S1). We performed smart-seq2 for full-length transcriptome profiling on the TV and LV cells. After stringent filtration, 265 individual TLV cells (128 TV cells and 137 LV cells) were retained for further analyses. As a result, these TV and LV cells were integrated and classified into six clusters using the function FindCluster in Seurat, while no distinct clusters were found between TV and LV (Figure 1B). To identify the properties of the different cell clusters, we identified and analyzed the differentially expressed genes (DEGs) in each cell cluster, including previously identified cell-type-specific markers. As shown in Figure 1C, the first cluster specifically expressed *COL1A1* [19] and *EBF2* [20], as well as osteoblast (OB) development-related genes such as *OGN* [21] and *GAS2* [22], implying that this cell cluster could be classified as OB. The second cluster specifically expressed *LUM* [23], *DCN* [24], and *TCF4* [25], suggesting that this cell cluster could be identified as fibroblast (FB). The third cluster expressed *HMGB2*, as well as cell mitosis-related genes, such as *TOP2A*, *MXD3*, *CDCA3*, *CDC20*, and *CKAP2.* As *HMGB2* plays a role in osteogenesis [26] and mesenchymal stem cell (MSC) differentiation [27], this cell cluster could be identified as stroma cell (SC). The fourth cluster *MATN1*, *COL11A1*, *COL11A2*, *MATN4*, and *MATN3* [28], as well as cartilage (CT) development-related genes, such as *CNMD*, *EPYC*, *HAPLN1*, and *PCOLCE2* [29], strongly suggesting that this cell cluster could be identified as CT. The fifth cluster expressed *CD248* [30] and *BMP4* [31], as well as MSC differentiation-related genes, such as *ASB9*, *MAB21L2*, *SERPINF1*, *NNAT*, and *CLDN11*, indicating that this cell cluster could be classified as MSC. While the sixth cluster expressed *CD34* [32, 33] and *CD93* [34], as well as angiogenesis-related genes, such as *PRCP* [35], *EMCN* [36], suggesting that this cell cluster contained hemogenic endothelial cell (HEC).

**Figure 1.**
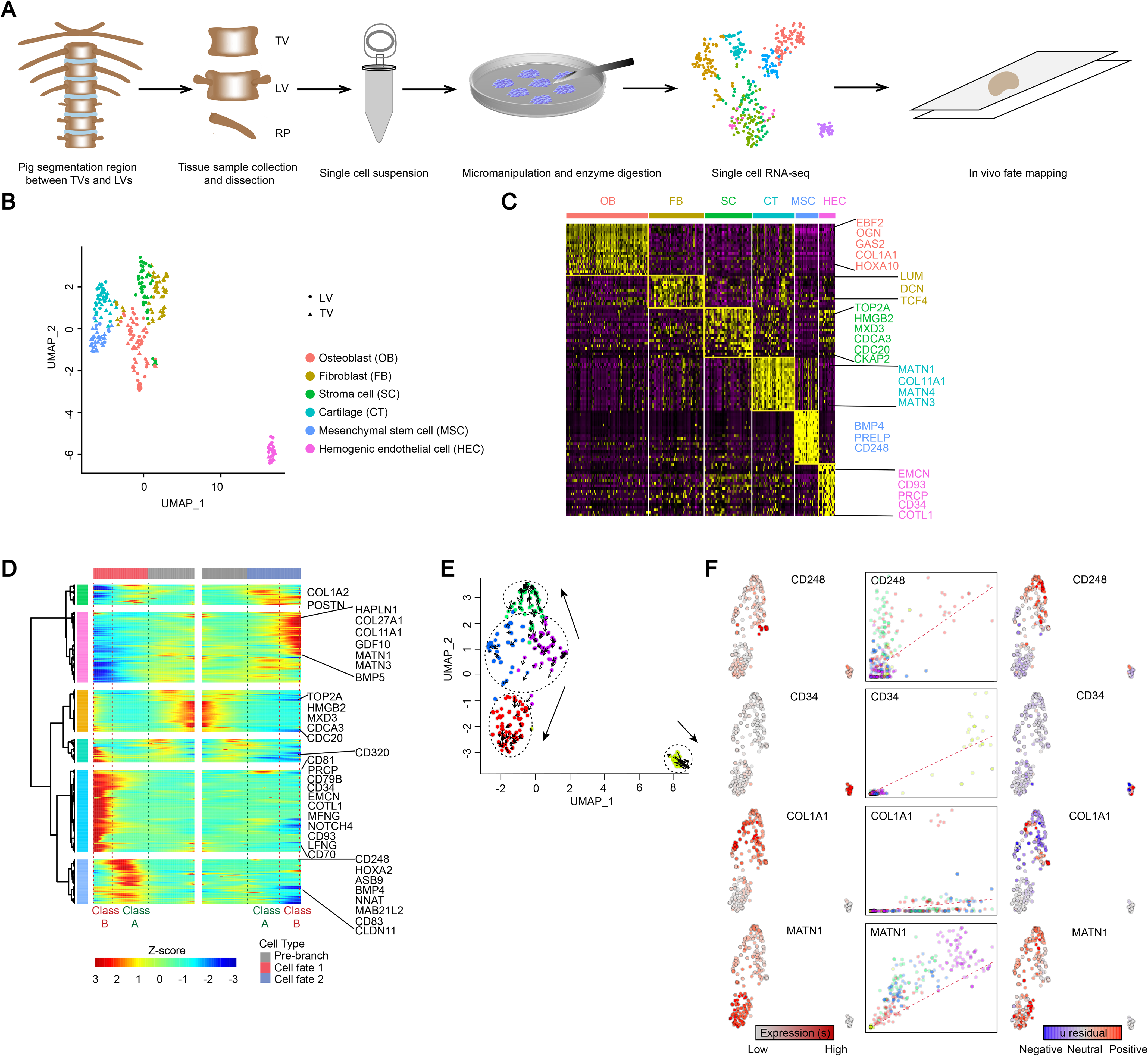
Characterization of TLV cell types and states in pigs. **A**. Overview of the experimental design. Segmentation regions between TV and LV dissected in pigs and dissociated into single-cell suspensions by micromanipulation and enzyme digestion. Smart-seq2 was used for single-cell RNA sequencing. **B**. Integrated UMAP plot of TV and LV cell clusters. LV and TV are indicated by dots and triangles, respectively. **C**. Gene expression patterns of each cell cluster in (B). Cell-type-enriched genes are listed on right and are labeled in the same colors as corresponding cell types. CT, cartilage; FB, fibroblast; OB, osteoblast; SC, stroma cell; HEC, Hemogenic endothelial cell; MSC, mesenchymal stem cell. **D**. Bifurcation of the 799 top TLV DEGs along two branches clustered hierarchically into five modules in a pseudo-temporal order. Development trajectories of TLV are shown on right and left, respectively. Class A and B indicate cell populations at earlier and later stages after segmentation between TV and LV, respectively; left, Class B, HEC, Class A, MSC; right, Class B, CT, Class A, OB. Representative genes are shown on the right. **E**. RNA velocity recapitulates dynamics of TLV cells differentiation. **F**. Expression pattern (left), unspliced-spliced phase portraits (center, cells colored according to the left), and u residuals (right) of TLV cells are shown for the repressed *CD248*, *CD34*, *COL1A1* and *MATN1* genes.

To reconstruct the developmental processes of TLV cells, we performed monocle-derived pseudo-time analysis. The TLV cells were successfully distributed along pseudo-temporal ordered paths consisting of a pre-branch and two cell fates: OB-CT (osteogenesis) and MSC-HEC (angiogenesis) (Figure 1D). These paths harbored cell-type-specific markers, including *COL1A2* [37], *MATN1* [38], *CD34* [32, 33], and *CD248* [30]. RNA velocity analysis further confirmed the general pattern of TLV cell differentiation associated with osteogenesis and angiogenesis (**Figure 1E**). The prediction of transcriptional dynamics of the developing pig TLV cells showed that OB and CT are from MSC, constituting the largest branch of differentiating lineages of the TLV cells. Our analysis revealed the expression of many marker genes, replicating the observation of expression of *CD248* [30] in the MSC zone, *COL1A1* [19] in the OB zone, *MATN1* [38] in the CT zone, and *CD34* [32,33] in the HEC zone (Figure 1F).

### Cell heterogeneity and *HOXA10* expressional difference between developing TV and LV

To explore the cell constitutional difference, we compared the fraction of cell populations in TV and LV. Cell cluster analysis revealed that both TV and LV contained six clusters of cells, replicating the observation with the whole cell samples (Figure S2), however, the fractions of cells in some clusters differ between TV and LV. The highest fractions were in CT cells from both groups, but showed no significant fraction difference (Permutation test, n = 1,000,000 replicates; P = 0.51). For the FB, SC, HEC and MSC cell clusters, the fraction difference was statistically significant when TV and LV cells were compared (FB, P=1.16×10^−2^; SC, P=2.25×10^−2^; HEC, P=7 ×10^−2^; MSC, P=5.04×10^−3^). While a difference for the OB cluster was observed, this fraction difference did not reach statistical significant (P=0.15).

Investigation of expressional marker in the cell clusters revealed differences between TV and LV. Comparison of the top 20 genes with the highest expression level in each cell cluster showed that most of the genes were shared by TV and LV cell clusters, however, there was a difference in the order of expression in TV and LV. Interestingly, we found that *HOXA10* showed differential gene expression levels in the OB cell clusters. We observed that *HOXA10* was the top gene with the highest expression level in OB from LV cells, but was absent from the top 20 genes in OB from TV cells. Additionally, *HOXA10* were not expressed in most TV and RP cells (118 of 128 TV cells and 174 of 199 RP cells, with FPKM < 1), while *HOXA10* had a high expression level in LV cells (79 of 137 LV cells, with FPKM > 5). Further validation using all of the single cell transcriptome data showed that the expression of *HOXA10* was largely restricted to cell clusters in LV, but was almost absent in TV cell clusters (Figure 2A). In the comparison of gene expression levels in OB from TV and LV cells, *HOXA10* showed the largest expressional bias toward LV (Figure 2B). *HOXA10* showed a wide expressional bias toward all LV cell clusters, in comparison to TV cell clusters. A similar, but incomplete pattern was observed for *HOXC10*, but not for *HOXD10* (Figure 2C). Combining all these data indicates that *HOXA10* may function as a determining factor that separate the sampled cells into either the TV or LV lineage.

**Figure 2.**
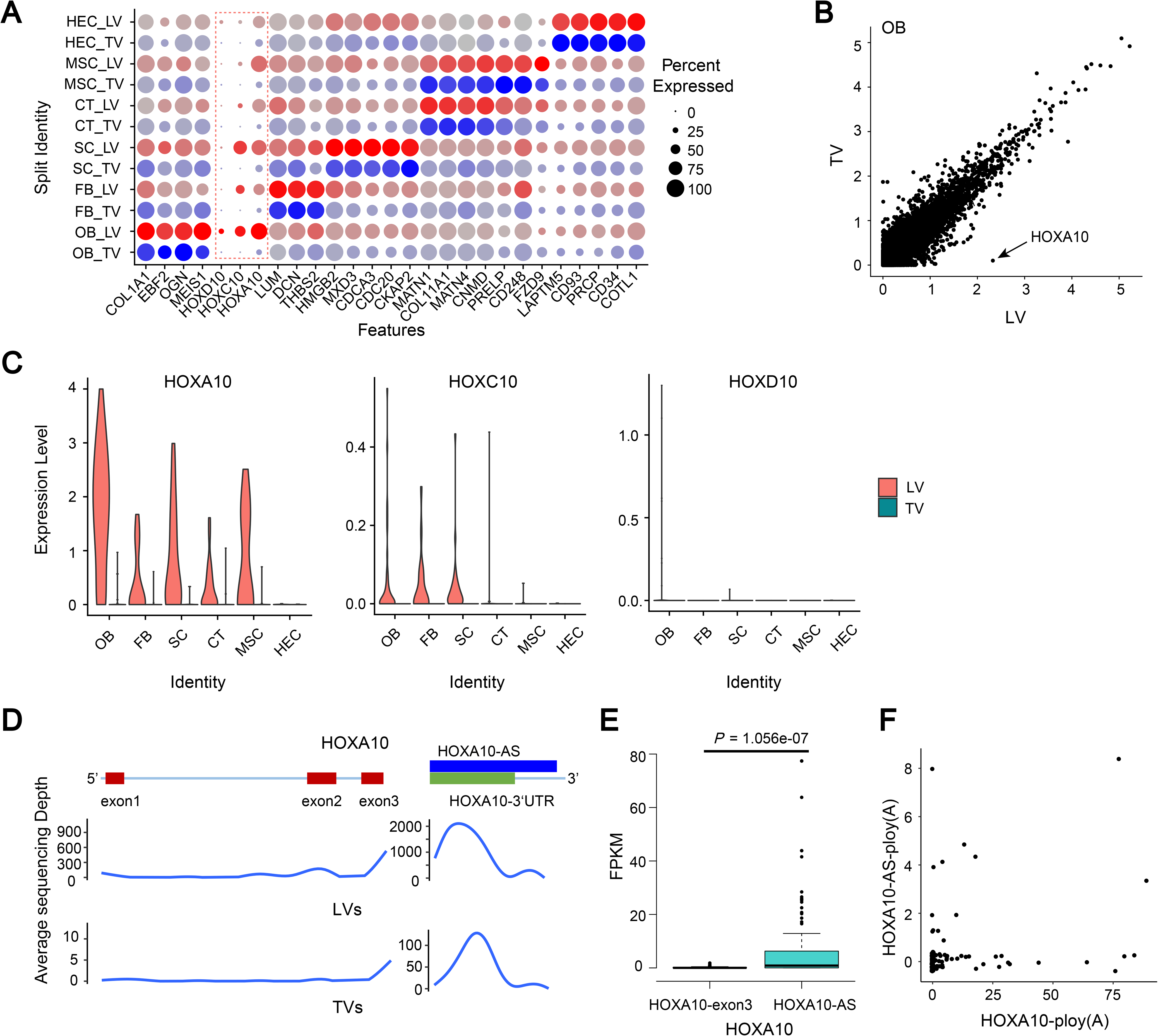
Characterization of cell-type-specific constitutional difference between TV and LV in pigs. **A**. Median scaled ln-normalized gene expression of selected DEGs for LV and TV cell clusters. FB, fibroblast; CT, cartilage; SC, stroma cell; OB, osteoblast; HEC, Hemogenic endothelial cell; MSC, mesenchymal stem cell. **B**. Scatter plot comparing the average expression of genes from both the LV and TV OB population. **C**. Vinplot comparing *HOXA10* expression of each cell cluster from both the LV and TV cells. **D**. Average sequencing depth of *HOXA10* exons and UTR and *HOXA10*-AS of 137 LV and 128 TV cells. Brown boxes represent exons of *HOXA10*; Green box represents 3’UTR of *HOXA10*; Blue box represents *HOXA10*-AS (*HOXA10*-antisense). **E**. Boxplot comparing the FPKM of exon3 and 3’-UTR of *HOXA10* from 137 LV cells. P value is from an unpaired two-sided Welch’s t-test with correction for multiple comparisons. **F**. Reads counts harboring *HOXA10*-ploy(A) and *HOXA10*-AS-ploy(A) at the *HOXA10*-3’UTR locus from 137 LV cells.

To further characterize the expression of *HOXA10* in developing TV and LV in pig embryo, we compared the distribution of sequencing reads at this loci (Figure 2D). On average, the sequencing depth of the reads in the *HOXA10* coding sequence was about 47.87X in LV cells and only 0.24X in TV cells. Unexpectedly, the sequencing depth in the 3’-UTR region of *HOXA10* in both TV (70.57X) and LV (1547.88X) were at least 30-fold higher than for the coding sequence. An analysis of the gene structure showed that an antisense RNA gene, *HOXA10* antisense (*HOXA10*-AS), overlaps the 3’-UTR on the opposite strand of the *HOXA10* gene. A possible explanation for the observed higher sequencing depth at the 3’-UTR is either a higher level of *HOXA10*-AS expression or *HOXA10* 3’-UTR expression, since single cell transcriptome sequencing cannot distinguish strand-specific RNA expression. Nevertheless, the average FPKM of *HOXA10*-exon3 were extremely lower than those for *HOXA10*-AS among the 137 LV cells (Figure 2E), with 0.18 for *HOXA10*-exon3 and 6.58 for HOXA10-AS (*p* = 1.056e-07, unpaired two-sided Welch’s t-test). To estimate the contribution of *HOXA10* and *HOXA10*-AS expression to the observed high levels of RNA sequencing depth for the *HOXA10* 3’-UTR genomic region, strand-specific expression was quantified using reads containing ploy(A) tail (Figure 2F). Among the 137 LV cells, *HOXA10* ploy (A) tail were detected in 44 cells, while *HOXA10*-AS tail were detected only in 13 cells. In addition, the number of reads containing *HOXA10* ploy (A) tail was also much higher than the number of reads containing *HOXA10*-AS ploy (A) tail, with an estimate of tenfold of the number. This implies that the high read sequencing depth from the *HOXA10* 3’-UTR genomic region were mainly due to *HOXA10* expression, rather than *HOXA10*-AS expression. We failed to find any reads containing *HOXA10* or *HOXA10*-AS poly (A) sequences in the 128 TV cells, possibly due to the low expression levels of this loci.

### Characterization of RP cell types and states

There is insufficient knowledge on the gene expression profiles involved in RP development in vertebrates. To understand this process, we collected 400 single cells from three RP close to the TLV segmentation joint for Smart-seq2 transcriptome profiling. After stringent filtration and classification by Seurat, 199 RP single cells were retained and classified into six clusters (Figure 3A and 3B). The first cluster expressed *TBX3* [39], *ASPN* [40], *YAP1* [41], and *SEMA3A* [42], suggesting that this cell cluster could be classified as FB. The second cluster expressed *MATN1*, *COL11A1*, *COL11A2*, *MATN4*, and *MATN3* [28,38], indicating that this cell cluster could be recognized as CT. The third cluster expressed *HMGB2*, *TOP2A*, *MXD3*, *CDCA3*, *CDC20*, and *CKAP2*, suggesting that this cell cluster contained SC [26,27]. The fourth cluster specifically expressed *EBF2*, *OGN*, *COL3A1*, and *COL1A1* [19,37], implying that this cell cluster could be identified as OB. The fifth cluster expressed *BMP4*, *FOS*, *FOSB*, *GADD45B*, and *CD248* [30,31], suggesting that this cell cluster could be classified as MSCs. The sixth cluster expressed *LAPTM5*, *PRCP*, *COTL1*, *CD93*, and *CD34* [32–35], indicating that this cell cluster resembled HEC.

**Figure 3.**
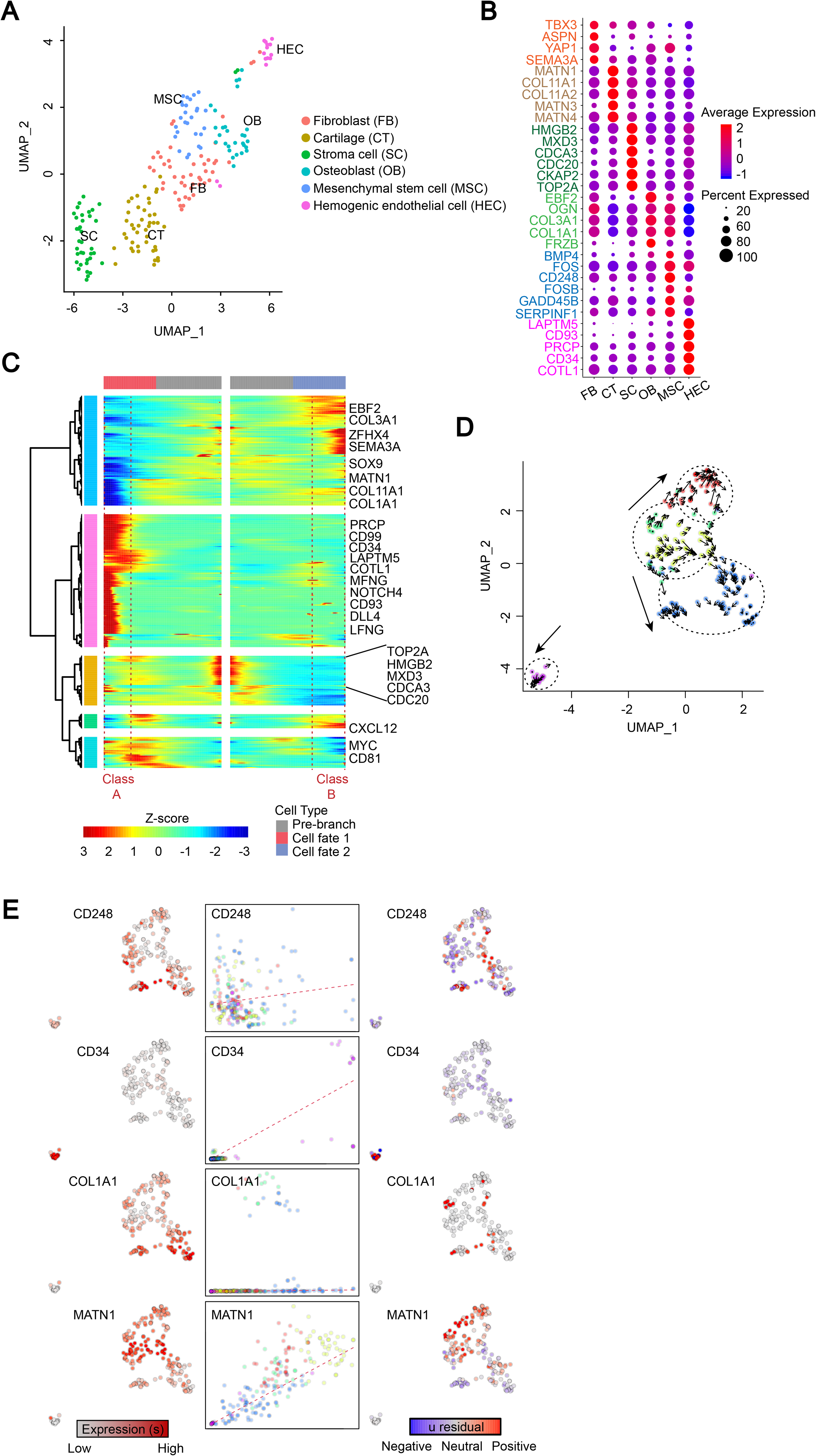
Characterization of RP cell types and states in pigs. **A**. UMAP plot of 199 RP cell clusters. FB, fibroblast; CT, cartilage; SC, stroma cell; OB, osteoblast; HEC, Hemogenic endothelial cell; MSC, mesenchymal stem cell. **B**. Median scaled ln-normalized gene expression of 31 selected DEGs for RP cell clusters from (A). Cell-type-enriched genes are listed on right and are labeled in the same colors as corresponding cell types. Gene expression level is simultaneously indicated by spot size and color intensity. **C**. Bifurcation of the 381 top RP DEGs along two branches clustered hierarchically into five modules in a pseudo-temporal order. Development trajectories of RP are shown on the right and left, respectively. Class A, HEC; Class B, OB. Gene expression level is indicated by color intensity. Representative genes are shown on the right. **D**. RNA velocity recapitulates the dynamics of the RP cells differentiation. **E**. Expression pattern (left), unspliced-spliced phase portraits (center, cells colored according to the left), and u residuals (right) of the RP cells are shown for the repressed *CD248*, *CD34*, *COL1A1* and *MATN1* genes.

Further, we performed monocle-derived pseudo-time analysis to reconstruct the RP developmental process. RP cells were distributed along a pseudo-temporally ordered paths consisting of a pre-branch and two cell fates, HEC (angiogenesis) and OB (osteogenesis) (Figure 3C). These paths harbored cell-type-specific markers, including *CD34* [32,33], *CD93* [34], *EBF2* [20], *COL1A1* [19], and *SOX9* [43], consistent with the gene expression patterns seen in Figure 3B. Similar results were obtained from a RNA velocity analysis that predicted the transcriptional dynamics of the developing pig RP cells compared to TLV cells (Figures 3D and 3E), indicating a robustness for the classification of angiogenesis and osteogenesis during the RP developmental process.

### Osteogenesis during TLV and RP development

To reveal the features of osteogenesis during TLV and RP development in pigs, we conducted a weighted gene co-expression network analysis (WGCNA) to construct a gene correlation network. As a CT marker and top transcriptional gene in the CT cluster, *MATN1* and its correlated genes were separated to build the osteogenic network. *MATN1* has been identified as a vital gene for cartilage networks in humans and mice [28,38]. Here, the hub genes correlated with *MATN1* included *COL11A1*, *COL11A1*, *COL2A1*, *EPYC*, *PCOLCE2*, *HAPLN1*, and *CNMD*, most of which are key genes involved in bone formation and remodeling [44] (Figure 4A).

**Figure 4.**
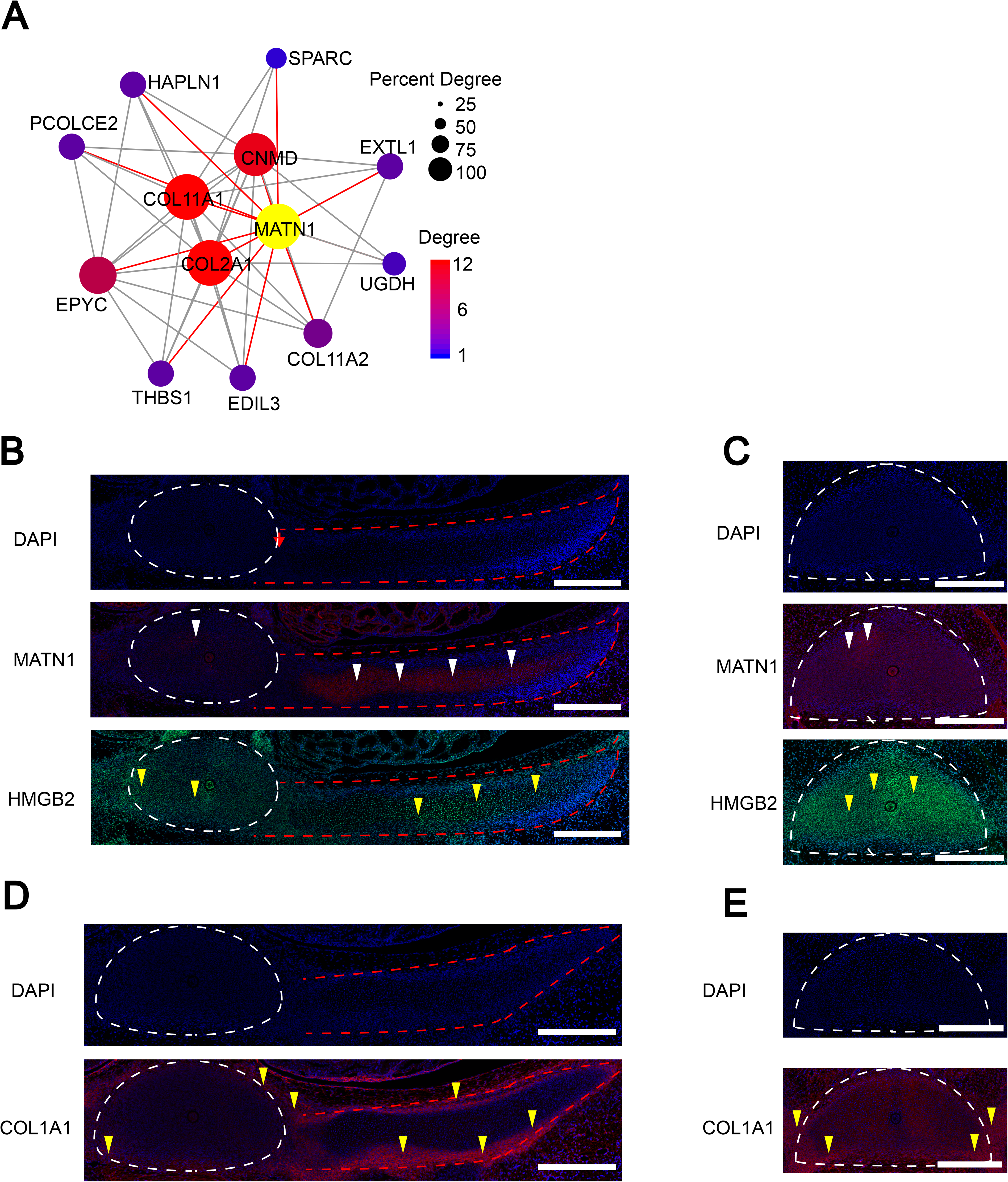
Osteogenesis during TLV and RP development in pigs. **A**. Module visualization of the network connections and associated functions using MATN1 as a hub gene. Gene connected intramodular degree is simultaneously indicated by spot size and color intensity. Hub gene MATN1 is indicated in yellow. **B**-**C**. Immunofluorescence analysis of MATN1 and HMGB2 in TV, RP, and LV. Gold and green indicate MATN1 and HMGB2, respectively. White triangles, MATN1+ cells; Yellow triangles, HMGB2+ cells. **B**. White and red dashed lines indicate regions of TV and RP (left and right), respectively. **C**. White dashed line indicates region of LV. Scale bar, 400 μm. **D**-**E**. Immunofluorescence analysis of COL1A1 in TV, RP, and LV. Gold indicates COL1A1. Yellow triangles, COL1A1+ cells. **D**. White and red dashed lines indicate regions of TV and RP (left and right), respectively. **E**. White dashed line indicates region of LV. Scale bar, 400 μm.

To confirm the spatial relationships among osteogenic cell types identified by Smart-seq2, we performed immunofluorescence imaging of TV and LV sections using pig embryo at 27 dpf. MATN1 [38], COL1A1 [19], and HMGB2 [26,27], which were identified as markers for CT, OB, and SC, respectively, were selected based on Gene Ontology (GO) analysis of DEGs for immunofluorescence analysis. Signals for the three proteins were detected in TLV and RP (Figure 4B-E). MATN1 is a secreted protein and was also detected inside the TLV and RP in our current study. In addition, HMGB2 was mainly detected in the nuclei and highly expressed in RP rather than in TLV, indicating that RP developed at an earlier stage of progenitor cell differentiation than TLV. COL1A1 is a secreted protein detected at the edge of TLV and RP. In terms of osteogenesis, our data imply that COL1A1 was first expressed at the edge of neonatal bone to generate OB, after which MATN1 is rapidly activated inside the neonatal bone to remodel and form CT during TLV and RP development in pigs at 27 dpf.

### Angiogenesis during TLV and RP development

Previous studies have shown that angiogenesis and osteogenesis are coupled in a specific vessel subtype during bone formation [36,45], while the features of angiogenesis during TLV and RP development remain unclear. As a HEC marker and top transcriptional gene in the HEC cluster, *CD34* and its related genes were separated to build an angiogenesis network by WGCNA [32, 33]. The hub genes correlated with *CD34* [32,33] include *CD93* [34], *PECAM1* (also known *CD31*) [36], *EMCN* [36], *PLVAP* [46], *F11R* (also known *CD321*) [47], *NPR1* [48] and *PRCP* [35], most of which are involved in angiogenesis (Figure 5A). These results indicated that *CD34* and *CD93* are two putative coordinators in the angiogenesis genetic network during early angiogenesis of TLV and RP development, in addition to *PECAM1* and *EMCN* [36]. Furthermore, pseudo-temporal order analysis of both TLV and RP based on the top DEGs suggested the involvement of Notch pathway components in angiogenesis, including *DLL4*, *MFNG*, *LFNG*, and *NOTCH4* (Figure 1D and 3C), consistent with previous reports suggesting that endothelial *Notch* activity promotes angiogenesis and osteogenesis in bone formation [45]. These results confirmed again that the sixth cluster of TLV and RP cells is HEC rather than a hematopoietic stem cell.

**Figure 5.**
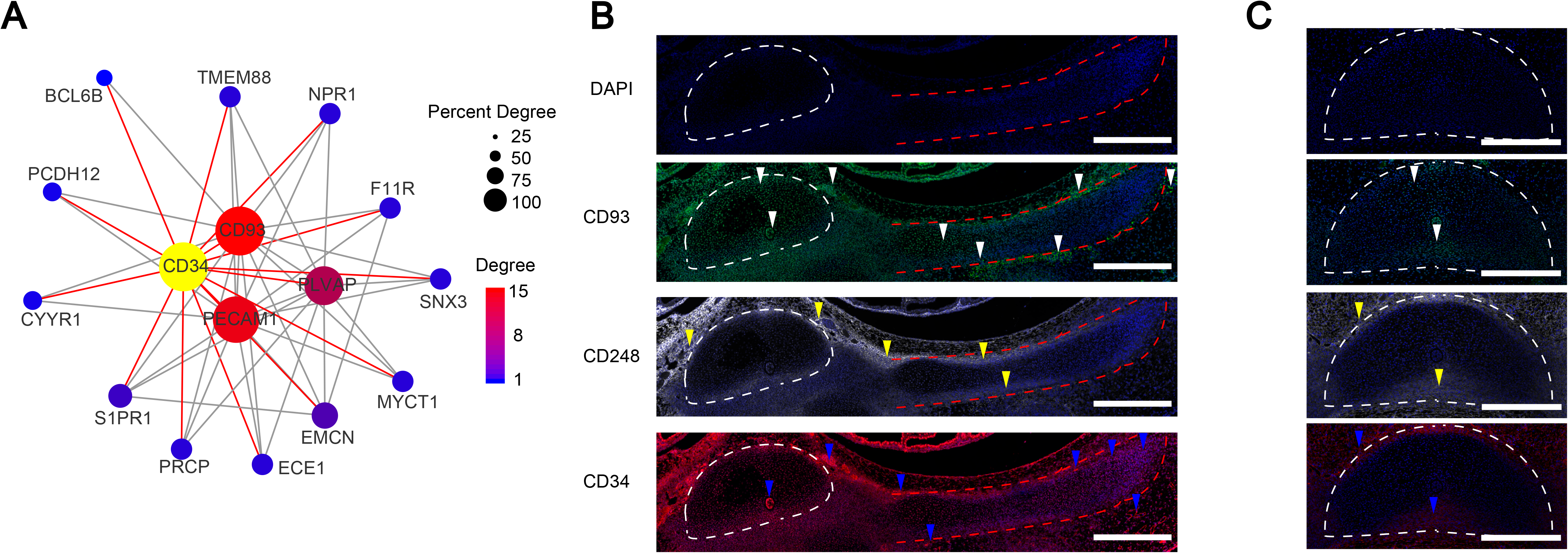
Angiogenesis during TLV and RP development in pigs. **A.** Module visualization of the network connections and associated functions using CD34 as a hub gene. Gene connected intramodular degree is simultaneously indicated by spot size and color intensity. Hub gene CD34 is indicated in yellow. **B**-**C**. Immunofluorescence analysis of CD93, CD248 and CD34 in TV, RP, and LV. Green, white and gold indicate CD93, CD248, and CD34, respectively. White triangles, CD93+ cells; Yellow triangles, CD248+ cells; Blue triangles, CD34+ cells. **B**. White and red dashed lines indicate region of TV and RP (left and right), respectively. **C**. White dashed line indicates region of LV. Scale bar, 400 μm.

To confirm the spatial relationships among the angiogenesis cell types identified by Smart-seq2, immunofluorescence imaging of TV and LV sections were performed using pig embryo at 27 dpf. CD34 [32,33], CD93 [34], and CD248 [30], which were identified as markers of HEC or MSC, were selected based on GO analysis of the DEGs for immunofluorescence analysis (Figure 5B and 5C). Protein signals of the three genes were detected in TLV, RP, and their surrounding tissues. CD34 is a secreted protein and was highly expressed in the notochord and tissues surrounding TLV and RP but was relatively less in TLV and RP. CD93 was expressed at a high level at the edge of RP, as well as in TLV and RP. In contrast, CD248, which is a membrane protein, was expressed at a high level at the edge of TLV and RP, but with almost no expression inside TLV or RP, similar to COL1A1, implying the synergistic action between MSC and OB during skeletal system development and remodeling. These data indicate that angiogenesis and osteogenesis are coupled by specific HESs during TLV and RP development in pigs at 27 dpf.

## Discussion

In this study, we demonstrated that the domestic pig can be a valuable animal model for exploring the molecular mechanisms underlying TLV transition and rib development in vertebrates. Despite the inter-specific difference of embryonic developmental processes, the overlap between the expressed genes observed in pigs and those reported in earlier mice models implies the presence of a conserved regulatory feature of TLV development among species [19,32,36]. The domestic pig may have advantages in exploring the genetic mechanisms underlying the TLV transition, since the number of TLV in different domestic pigs vary [17]. The intra-specific developmental variation may offer an opportunity for future studies to explore the genomic coordination that determines the TV and LV body identities, giving a low level of genomic noise within a single species. The results obtained in this study open a new window and provided valuable resources to expand such studies on TLV transition using pigs.

Our analysis revealed that the cell types in developmental TLV can be functionally clustered into two groups, corresponding to the osteogenesis (OB-CT) and angiogenesis (MSC-HEC) biological processes. RP cells were functionally correlated to osteogenesis (OB) and angiogenesis (HEC). Although this study may have been affected by the limited number of cells sampled for sc-RNA sequencing, our observations revealed a coupled process of angiogenesis and osteogenesis during TLV and RP development, highly consistent with the observations from previous studies during bone formation [36,45]. This implies that the number of sampled cells has substantial information for representing a cell atlas of developing TLV and RP. This study allows discovery of the temporal resolution of transcriptome kinetics using scRNA-seq data and provides fundamental insight relevant to abnormal TLV transition and RP development in vertebrates.

Previous studies via intervention of gene expression, whole mount in situ hybridization, and immunofluorescence analysis revealed a crucial role of *HOXA10* in TLV transition, while the molecular mechanism underlying *HOXA10* determining TLV transition remains elusive [7–9]. An important merit of this study is the discovery of the restricted high expression pattern of *HOXA10* to only OB of LV cells rather than in TV cells using our scRNA-seq data. In-depth dissection of the reads distribution revealed that most reads were restricted to the 3’ UTR of *HOXA10* rather than the coding region and overlapped with *HOXA10*-AS. *HOXA10*-AS is capable of repressing *HOXA10* sense expression [49,50]. The strand-specific expression analysis using reads containing ploy(A) tail revealed that the high read sequencing depth from *HOXA10* 3’-UTR genomic region were mainly due to *HOXA10* expression, rather than *HOXA10*-AS expression. These observations suggested that the TLV transition involves a putative expressional balance between the *HOXA10* and *HOXA10-AS* genes. The large amount of reads clustered in the *HOXA10* 3’-UTR genomic regions indicates the presence of a regulatory mechanism that blocks the expression of *HOXA10*-*AS* through an expression of a short transcript from the 3’-end of the *HOXA10* sense strand that is complementary to the *HOXA10*-*AS* sequence, implying an anti-*HOXA10*-AS role of the high levels of *HOXA10* 3’-UTR sequence within OB of LV cells.

It has been clearly shown that HEC are from the aorta-gonad-mesonephros region where hematopoiesis takes place and then migrate to the liver and bone marrow [51,52]. Previous studies on hematopoiesis focused on the liver and bone marrow, but insufficiently in somite and TLV and RP [53–55]. In zebrafish, somatic cells in embryos can trans-differentiate into hematopoietic cells showing that somite is an additional embryonic hematopoietic site validated by transgenic lineage tracing [56,57]. Our results revealed a specific vessel subtype HEC with distinct molecular and functional properties during TLV and RP development in pigs at 27 dpf. The angiogenesis gene network was established as early as four post-fertilization weeks in TLV and RP of pigs.

Bone developmental processes include four stages, i.e., pre-CT stage (spongy bone), CT stage, CT erosion (cancellous bone), and bone deposition (compact bone) [58]. Here, we focused on the stage during which spongy bone turns into CT but found no notable differences in the cell clusters between TV and LV. Sampling at 17 to 27 dpf may provide insights on these possible differences; however, it would be difficult to confirm the joint TLV partition as RP are not developed in pigs before 27 dpf [Li et al. unpublished data].

In summary, our comprehensive TLV and RP atlas at 27 dpf provides a valuable resource for understanding molecular programs and development temporary states during TLV transition and rib-genesis in vertebrates. Our approach using single-cell transcriptomics to study TLV and RP development in pigs provides a framework that could be applied to study temporal processes using other animal models.

## Materials and methods

### Sample collection and preparation of single-cell suspensions

TV, LV, and RP cells from three vertebrae close to the TLV segmentation joint of one LW pig embryo at 27 dpf were collected by micromanipulation and enzyme digestion. TV, LV, and RP samples were uniformly dissected into millimeter-sized pieces in Dulbecco’s phosphate-buffered saline (DPBS), supplemented with 10% fetal calf serum, transferred to tubes containing 1 mL of collagenase II (5 mg/ml) and dispase (2.5 mg/mL), and then incubated at 37 °C for 3–5 min. Digested tissue pieces were then filtered through a 40-mm nylon cell strainer (352340, BD Falcon), with the filtered cell suspension centrifuged at 500 g for 10 min at 37 °C. After removal of the supernatant, the cell pellets were washed using 1 × DPBS three times to remove fragments and then resuspended in Dulbecco’s Modified Eagle’s Medium (11995040, Gibco).

### Full-length single-cell RNA-seq library preparation and sequencing

We used the Smart-seq2 protocol for full-length single-cell RNA-seq according to the manufacturer’s instructions [59]. Briefly, single cells were transferred to lysis buffer with RNase Inhibitor in a 0.2-mL polymerase chain reaction (PCR) tube by mouth pipetting. First-strand cDNA synthesis was performed using a 25-bp oligo(dT)_30_VN primer for 3’ amplification. Second-strand cDNA was generated via PCR amplification. The newly synthesized double-stranded cDNA was then annealed to an index primer. Subsequently, the synthetic DNA was fragmented into 350-bp pieces by a Bioruptor® Sonication System (UCD300, Diagenode Inc.), and the reactions were purified by incubation with Ampure XP beads (A63880, Beckman) at room temperature for 5 min. After quality inspection using an Agilent 2100 High Sensitivity DNA Assay Kit (5067-4626, Agilent Technologies) based on the manufacturer’s instructions, sequencing was performed on an Illumina HiSeq 2000 platform using 150-bp paired-end sequencing via the Smart-seq2 protocol.

### Processing of single-cell RNA-seq data

Clean reads were aligned to the reference pig genome (genome build: Sscrofa 11.1) using Hisat2 (v2.0.5). We calculated and partitioned the uniquely mapped reads using StringTie (v1.3.5) and Ballgown (v2.16.0) [60]. The transcript counts of each cell were normalized to reads per kilobase of exon model per million mapped reads (FPKM). Overall, 800 individual cells were collected for single-cell cDNA amplification and 465 cells passed the quality control criteria. On average, there were 22 million mapped reads and 7,253 detected genes for each cell.

### Identification of cell type using UMAP

The Seurat (v3.0), dplyr (v0.7.0), and umap (v0.2.3.1) packages in R were applied to classify the 465 single cells into major cell types [61,62]. Only single cells with a gene expression number > 2,000 were considered, and only genes with normalized expression levels greater than one (1) expressed in at least three single cells were retained. Finally, we obtained 22,517 genes across the 465 samples for clustering analysis. We conducted principal component analysis (PCA) of the genes from the remaining cells using the FindVariableFeatures function (selection.method = “vst”, nfeatures = 2000)”. The principal components (PCs) used to partition cells were selected using Jackstraw (100 replicates). The first 10 PCs were used to perform UMAP based on the RunUMAP and FindClusters functions. We obtained six cell clusters in both TLV and RP.

### Identification of cell-type-specific expressed genes

Genes that were differentially expressed in each cluster were identified using the function ‘FindAllMarkers’ in Seurat against the normalized gene expression data and were then tested by ‘roc’ and DESeq2 [59]. Here, both min.pct and thresh.use values greater than 0.25 were selected as the cut-off for gene selection. Samtools and bedtools were used to calculate sequencing depth and reads harboring strand-specific poly (A) tail of each TLV cell [63,64]. The Database for Annotation, Visualization, and Integrated Discovery (DAVID Gene Functional Classification Tool) Bioinformatics Resource was used for functional annotation and GO analysis [65].

### Pseudo-time analysis

The Monocle2 package was used to analyze pseudo-time trajectories to determine developmental transitions in RP and the segmentation processes between TV and LV [66]. We used cell-type-specific expressed genes identified by Seurat to sort cells into pseudo-time order. ‘DDRTree’ was applied to reduce the dimensions and the visualization functions ‘plot_cell_trajectory’, ‘plot_genes_branched_pseudotime’, and ‘plot_genes_branched_heatmap’ were used to display the pseudo-time gene branched trajectories, pseudo-times, and heatmaps, respectively.

### RNA velocity analysis

Read annotations for the Smart-seq2 output data was performed using the velocyto.py command-line tools according to the manual [67]. Genome annotations Sscrofa11.1 and Sscrofa11.1.101 from Ensembl were used to count, and sort reads into three categories: ‘spliced’, ‘unspliced’ or ‘ambiguous’. The loom file generated was loaded into velocyto.R. Finally, coordinates from the Seurat's UMAP analysis were used to embed the velocity results.

### WGCNA and gene correlation network construction

WGCNA was performed on the normalized gene expression data measured in FPKM, using unsigned correlation, soft-threshold power of six, and minimum module size of 120 members [68]. We then generated an independent list of hub genes (kME > 0.9) for each skeletal region. Finally, the co-expression gene network was visualized using VisANT and Cytoscape [69, 70]. The Benjamini-Hochberg method was used to correct for multiple testing when calculating the significance of the correlations among modules and skeletal regions.

### Immunofluorescence staining analysis

Pig embryo at 27 dpf was fixed overnight in 10% neutral formalin fixed solution at room temperature. Thin 5-μm TV and LV paraffin-embedded sections were collected for immunofluorescence staining. Cell nuclei were stained using DAPI (Life Technologies, USA).

Primary antibodies used were: MATN1 (rabbit, 1:200, Biorbyt, orb94279), HMGB2 (Rabbit, 1:500, Abcam, ab67282), COL1A1 (rabbit, 1:200, Abcam, ab34710), CD34 (rabbit, 1:200, Biorbyt, orb348961), CD93 (rabbit, 1:200, Abcam, ab198854), and CD248 (mouse, 1:50, Santa Cruz Biotechnology, sc-377221). Secondary antibody used was anti-rabbit IgG (1:500, ZSGB-BIO, ZDR-5003).

### Image analysis and data processing

Images of the paraffin sections were collected by digitizing the images using a Leica Aperio Versa 200 slide scanner (Germany). All images were processed using ImageScope.

### Ethical statement

Pig embryonic and fetal sample collection and single-cell transcriptome study were carried out under the approval of the Kunming Institute of Zoology, Chinese Academy of Sciences, China (SMKX-20191213-01). All experiments were done following the International Review Board and Institutional Animal Care and Use Committee guidelines.

This manuscript has not been published or presented elsewhere in part or in entirety, and is not under consideration by another journal. There are no conflicts of interest to declare.

## Data availability

Single-cell RNA-seq data used in this study were deposited into the Genome Sequence Archive of the Beijing Institute of Genomics, Chinese Academy of Sciences (http://gsa.big.ac.cn/) under accession number CRA002562.

## CRediT author statement

**Jianbo Li:** Formal analysis, Writing-original draft, Writing-review & editing, Visualization. **Ligang Wang:** Formal analysis, Writing-original draft. **Dawei Yu:** Collect the embryo and single cells. **Junfeng Hao:** Conduct the immunofluorescence analysis. **Longchao Zhang:** Supply the pregnant LW pig. **Adeniyi C. Adeola:** Improve the manuscripts. **Bingyu Mao:** Revise the manuscripts. **Yun Gao:** Resources. **Shifang Wu:** Resources. **Chunling Zhu:** Resources. **Yongqing Zhang:** Revise the manuscripts. **Jilong Ren:** Resources. **Changgai Mu:** Resources. **David M. Irwin:** Improve the manuscripts. **Lixian Wang:** Project administration, Funding acquisition. **Tang Hai:** Project administration, Funding acquisition. **Haibing Xie:** Formal analysis, Writing-original draft. **Yaping Zhang:** Supervision, Project administration, Funding acquisition.

## Acknowledgments

The project was supported by the Strategic Pioneer Program of the Chinese Academy of Sciences (XDA24010107), Ministry of Agriculture of China (Grant No. 2016ZX08009003-006), China Agriculture Research System (CARS-35), and Agricultural Science and Technology Innovation Project (ASTIP-IAS02). This work was supported by the Animal Branch of the Germplasm Bank of Wild Species, Chinese Academy of Sciences (the Large Research Infrastructure Funding).

## Supplementary material

**Figure S1 Pig TLV and RP separation by micromanipulation and enzymatic digestion at 27 dpf (left to right)** Scale bar, 2 mm.

**Figure S2 Characterization of cellular populations of TVs and LVs in pigs** UMAP plot of 137 LV (A) and 128 TV (C) cell clusters, respectively. Gene expression patterns of each cell cluster in LV (B) and TV (D), respectively. Cell-type-enriched genes are listed on the right and are labeled in the same colors as their corresponding cell type. CT, cartilage; SC, stroma cell; OB, osteoblast; FB, fibroblast; HEC, hematopoietic stem cell; MSC, mesenchymal stem cell.

## References

[1] Gilbert SF, Barresi MJ. Development biology, 11th ed. 2016.

[2] Gomez C, Özbudak EM, Wunderlich J, Baumann D, Lewis J, Pourquié O. Control of segment number in vertebrate embryos. Nature 2008; 454: 335–339.

[3] Wilson V, Olivera-Martinez I, Storey KG. Stem cells, signals and vertebrate body axis extension. Development 2009; 136: 1591–1604.

[4] Woltering JM. From Lizard to Snake; Behind the Evolution of an Extreme Body Plan. Curr Genomics 2012; 13: 289–299.

[5] Jurberg AD, Aires R, Varela-Lasheras I, Novoa A, Mallo M. Switching axial progenitors from producing trunk to tail tissues in vertebrate embryos. Dev Cell 2013; 25: 451–462.

[6] Aires R, Jurberg AD, Leal F, Novoa A, Cohn MJ, Mallo M. Oct4 Is a Key Regulator of Vertebrate Trunk Length Diversity. Dev Cell 2016; 38: 262–274.

[7] Wellik DM, Capecchi MR. Hox10 and Hox11 Genes Are Required to Globally Pattern the Mammalian Skeleton. Science 2003; 301: 63–367.

[8] Guerreiro I, Nunes A, Woltering JM, Casaca A, Novoa A, Vinagre T, et al. Role of a polymorphism in a Hox/Pax-responsive enhancer in the evolution of the vertebrate spine. Proc Natl Acad Sci USA 2013; 110: 10682–10686.

[9] McIntyre DC, Rakshit S, Yallowitz AR, Loken L, Jeannotte L, Capecchi MR, et al. Hox patterning of the vertebrate rib cage. Development 2007; 134: 2981–2989.

[10] Mu Q, Chen Y, Wang J. Deciphering Brain Complexity Using Single-cell Sequencing. Genom Proteom Bioinf 2019; 17: 344–366.

[11] Xhangolli I, Dura B, Lee G, Kim D, Xiao Y, Fan R. Single-cell Analysis of CAR-T Cell Activation Reveals A Mixed TH1/TH2 Response Independent of Differentiation. Genom Proteom Bioinf 2019; 17: 129–139.

[12] Coordination O, Coordination L, Preparation L, Annotation CT, Investigators P. Single-cell transcriptomics of 20 mouse organs creates a Tabula Muris. Nature 2018; 562: 367–372

[13] Camp JG, Platt RJ, Treutlein B. Mapping human cell phenotypes to genotypes with single-cell genomics. Science 2019; 365: 1401–1405.

[14] Han X, Wang R, Zhou Y, Fei L, Sun H, Lai S, et al. Mapping the Mouse Cell Atlas by Microwell-Seq. Cell 2018; 172: 1091–1107.e1017.

[15] Han X, Zhou Z, Fei L, Sun H, Wang R, Chen Y, et al. Construction of a human cell landscape at single-cell level. Nature 2020; 562: 367–372.

[16] Baryawno N, Przybylski D, Kowalczyk MS, Kfoury Y, Severe N, Gustafsson K, et al. A Cellular Taxonomy of the Bone Marrow Stroma in Homeostasis and Leukemia. Cell 2019; 177: 1915–1932.e1916.

[17] Borchers N, Reinsch N, Kalm E. The number of ribs and vertebrae in a Piétrain cross: variation, heritability and effects on performance traits. J Anim Breed Genet 2004; 121: 392–403.

[18] Zhang Z, Sun Y, Du W, He S, Liu M, Tian C. Effects of Vertebral Number Variations on Carcass Traits and Genotyping of VRTN Candidate Gene in Kazakh Sheep. Asian-Austral J Anim 2017; 30: 1234–1238.

[19] Dacic S, Kalajzic I, Visnjic D, Lichtler A, Rowe D. Col1a1[driven transgenic markers of osteoblast lineage progression. J Bone Miner Res 2001; 16: 1228–1236.

[20] Kieslinger M, Folberth S, Dobreva G, Dorn T, Croci L, Erben R, et al. EBF2 regulates osteoblast-dependent differentiation of osteoclasts. Dev Cell 2005; 9: 757–767.

[21] Rehn AP, Cerny R, Sugars RV, Kaukua N, Wendel M. Osteoadherin is upregulated by mature osteoblasts and enhances their in vitro differentiation and mineralization. Calcified Tissue Int 2008; 82: 454–464.

[22] Capulli M, Rufo A, Teti A, Rucci N. Global transcriptome analysis in mouse calvarial osteoblasts highlights sets of genes regulated by modeled microgravity and identifies a “mechanoresponsive osteoblast gene signature”. J Cell Biochem 2009; 107: 240–252.

[23] Pilling D, Vakil V, Cox N, Gomer RH. TNF-α–stimulated fibroblasts secrete lumican to promote fibrocyte differentiation. Proc Natl Acad Sci USA 2015; 112: 11929–11934.

[24] Ferdous Z, Peterson SB, Tseng H, Anderson DK, Iozzo RV, Grande-Allen KJ. A role for decorin in controlling proliferation, adhesion, and migration of murine embryonic fibroblasts. J Biomed Mater Res A 2010; 93: 419–428.

[25] Contreras O, Rebolledo D, Oyarzún J, Brandan E. Fibroblasts Tcf4 and mesenchymal progenitors PDGFRα correspond to the same cell type and are increased in skeletal muscle dystrophy, denervation and chronic damage. F1000Research 2019; 8.

[26] Taniguchi N, Caramés B, Ronfani L, Ulmer U, Komiya S, Bianchi ME, et al. Aging-related loss of the chromatin protein HMGB2 in articular cartilage is linked to reduced cellularity and osteoarthritis. Proc Natl Acad Sci USA 2009; 106: 1181–1186.

[27] Taniguchi N, Caramés B, Hsu E, Cherqui S, Kawakami Y, Lotz M. Expression patterns and function of chromatin protein HMGB2 during mesenchymal stem cell differentiation. J Biol Chem 2011; 286: 41489–41498.

[28] Pei M, Luo J, Chen Q. Enhancing and maintaining chondrogenesis of synovial fibroblasts by cartilage extracellular matrix protein matrilins. Osteoarthr Cartilage 2008; 16: 1110–1117.

[29] Miclea R, Siebelt M, Finos L, Goeman J, Löwik C, Oostdijk W, et al. Inhibition of Gsk3β in cartilage induces osteoarthritic features through activation of the canonical Wnt signaling pathway. Osteoarthr Cartilage 2011; 19: 1363–1372.

[30] Naylor AJ, Azzam E, Smith S, Croft A, Poyser C, Duffield JS, et al. The mesenchymal stem cell marker CD248 endosialin is a negative regulator of bone formation in mice. Arthritis & Rheumatism 2012; 64: 3334–3343.

[31] Li N, Pan S, Zhu H, Mu H, Liu W, Hua J. BMP4 promotes SSEA◻1+ hUC◻MSC differentiation into male germ◻like cells in vitro. Cell Proliferat 2014; 47: 299–309.

[32] Asahara T, Murohara T, Sullivan A, Silver M, van der Zee R, Li T, et al. Isolation of putative progenitor endothelial cells for angiogenesis. Science 1997; 275: 964–967.

[33] Sidney LE, Branch MJ, Dunphy SE, Dua HS, Hopkinson A. Concise review: evidence for CD34 as a common marker for diverse progenitors. Stem Cells 2014; 32: 1380–1389.

[34] Galvagni F, Nardi F, Maida M, Bernardini G, Vannuccini S, Petraglia F, et al. CD93 and dystroglycan cooperation in human endothelial cell adhesion and migration adhesion and migration. Oncotarget 2016; 7: 10090–10103.

[35] Hagedorn M. PRCP regulates angiogenesis in vivo. Blood 2013; 122: 1337–1338.

[36] Kusumbe AP, Ramasamy SK, Adams RH. Coupling of angiogenesis and osteogenesis by a specific vessel subtype in bone. Nature 2014; 507: 323–328.

[37] Yano H, Hamanaka R, Nakamura-Ota M, Adachi S, Zhang JJ, Matsuo N, et al. Sp7/Osterix induces the mouse pro-α2 I collagen gene Col1a2 expression via the proximal promoter in osteoblastic cells. Biochem Bioph Res Co 2014; 452: 531–536.

[38] Nicolae C, Ko Y-P, Miosge N, Niehoff A, Studer D, Enggist L, et al. Abnormal collagen fibrils in cartilage of matrilin-1/matrilin-3-deficient mice. J Biol Chem 2007; 282: 22163–22175.

[39] Han J, Yuan P, Yang H, Zhang J, Soh BS, Li P, et al. Tbx3 improves the germ-line competency of induced pluripotent stem cells. Nature 2010; 463: 1096–1100.

[40] Orr B, Riddick A, Stewart G, Anderson R, Franco O, Hayward S, et al. Identification of stromally expressed molecules in the prostate by tag-profiling of cancer-associated fibroblasts, normal fibroblasts and fetal prostate. Oncogene 2012; 31: 1130–1142.

[41] Piersma B, de Rond S, Werker PM, Boo S, Hinz B, van Beuge MM, et al. YAP1 is a driver of myofibroblast differentiation in normal and diseased fibroblasts. Am J Pathol 2015; 185: 3326–3337.

[42] Fukamachi S, Bito T, Shiraishi N, Kobayashi M, Kabashima K, Nakamura M, et al. Modulation of semaphorin 3A expression by calcium concentration and histamine in human keratinocytes and fibroblasts. J Dermatol Sci 2011; 61: 118–123.

[43] Zhou G, Zheng Q, Engin F, Munivez E, Chen Y, Sebald E, et al. Dominance of SOX9 function over RUNX2 during skeletogenesis. Proc Natl Acad Sci USA 2006; 103: 19004–19009.

[44] van Gool S, Emons J, Leijten J, Decker E, Yu X, Sticht C. Human fetal mesenchymal stem cells differentiating towards chondrocytes display a similar gene expression profile as growth plate cartilage. Regulation And Modulation Of Growth 2010; 135.

[45] Ramasamy SK, Kusumbe AP, Wang L, Adams RH. Endothelial Notch activity promotes angiogenesis and osteogenesis in bone. Nature 2014; 507: 376–380.

[46] Guo L, Zhang H, Hou Y, Wei T, Liu J. Plasmalemma vesicle-associated protein: A crucial component of vascular homeostasis. Exp Ther Med 2016; 12: 1639–1644.

[47] Fukuhara T, Kim J, Hokaiwado S, Nawa M, Okamoto H, Kogiso T, et al. A novel immunotoxin reveals a new role for CD321 in endothelial cells. PLoS One 2017; 12: e0181502.

[48] Yue B, Ma JF, Yao G, Yang MD, Cheng H, Liu GY. Knockdown of neuropilin-1 suppresses invasion, angiogenesis, and increases the chemosensitivity to doxorubicin in osteosarcoma cells - an in vitro study. European review for medical and pharmacological sciences 2014; 18: 1735–1741.

[49] Bagot CN, Troy PJ, Taylor HS. Alteration of maternal Hoxa10 expression by in vivo gene transfection affects implantation. Gene Ther 2000; 7: 1378–1384.

[50] Daftary GS, Troy PJ, Bagot CN, Young SL, Taylor HS. Direct regulation of beta3-integrin subunit gene expression by HOXA10 in endometrial cells. Mol Endocrinol 2002; 16: 571–579.

[51] Eilken HM, Adams RH. Dynamics of endothelial cell behavior in sprouting angiogenesis. Curr Opin Cell Biol 2010; 22: 617–625.

[52] Lancrin C, Sroczynska P, Serrano A, Gandillet A, Ferreras C, Kouskoff V, et al. Blood cell generation from the hemangioblast. J Mol Med Berl 2010; 88: 167–172.

[53] Morrison SJ, Scadden DT. The bone marrow niche for haematopoietic stem cells. Nature 2014; 505: 327–334.

[54] Popescu DM, Botting RA, Stephenson E, Green K, Webb S, Jardine L, et al. Decoding human fetal liver haematopoiesis. Nature 2019; 574: 365–371.

[55] Bian Z, Gong Y, Huang T, Lee CZW, Bian L, Bai Z, et al. Deciphering human macrophage development at single-cell resolution. Nature 2020; 582: 571–576.

[56] Nguyen PD, Hollway GE, Sonntag C, Miles LB, Hall TE, Berger S, et al. Haematopoietic stem cell induction by somite-derived endothelial cells controlled by meox1. Nature 2014; 512: 314–318.

[57] Qiu J, Fan X, Wang Y, Jin H, Song Y, Han Y, et al. Embryonic hematopoiesis in vertebrate somites gives rise to definitive hematopoietic stem cells. J Mol Cell Biol 2016; 8: 288–301.

[58] Olsen BR, Reginato AM, Wang W. Bone development. Annu Rev Cell Dev Biol 2000; 16: 191–220.

[59] Picelli S, Faridani OR, Björklund ÅK, Winberg G, Sagasser S, Sandberg R. Full-length RNA-seq from single cells using Smart-seq2. Nat Protoc 2014; 9: 171.

[60] Pertea M, Kim D, Pertea GM, Leek JT, Salzberg SL. Transcript-level expression analysis of RNA-seq experiments with HISAT, StringTie and Ballgown. Nat Protoc 2016; 11: 1650–1667.

[61] Satija R, Farrell JA, Gennert D, Schier AF, Regev A. Spatial reconstruction of single-cell gene expression data. Nat Biotechnol 2015; 33: 495–502.

[62] Butler A, Hoffman P, Smibert P, Papalexi E, Satija R. Integrating single-cell transcriptomic data across different conditions, technologies, and species. Nat Biotechnol 2018; 36: 411–420.

[63] Li H, Handsaker B, Wysoker A, Fennell T, Ruan J, Homer N, et al. The Sequence Alignment/Map format and SAMtools. Bioinformatics 2009; 25: 2078–2079.

[64] Quinlan AR. BEDTools: the Swiss-army tool for genome feature analysis. Current Protocols In Human Genetics 2014; 47.

[65] Huang DW, Sherman BT, Tan Q, Collins JR, Alvord WG, Roayaei J, et al. The DAVID Gene Functional Classification Tool: a novel biological module-centric algorithm to functionally analyze large gene lists. Genome Biol 2007; 8: R183.

[66] Qiu X, Mao Q, Tang Y, Wang L, Chawla R, Pliner HA, et al. Reversed graph embedding resolves complex single-cell trajectories. Nat Methods 2017; 14: 979.

[67] La Manno G, Soldatov R, Zeisel A, Braun E, Hochgerner H, Petukhov V, et al. RNA velocity of single cells. Nature 2018; 560: 494–498.

[68] Langfelder P, Horvath S. WGCNA: an R package for weighted correlation network analysis. BMC Bioinformatics 2008; 9: 559.

[69] Hu Z, Mellor JC, Wu J, Delisi C. VisANT: an online visualization and analysis tool for biological interaction data. BMC Bioinformatics 2004; 5: 17–17.

[70] Kohl M, Wiese S, Warscheid B. Cytoscape: Software for Visualization and Analysis of Biological Networks. Methods Mol Biol 2011; 696: 291–303.

