## Supplementary figures and images for "Single-cell RNA-sequencing reveals thoracolumbar vertebra heterogeneity and rib-genesis in pigs"

### Supplemental Figure 1

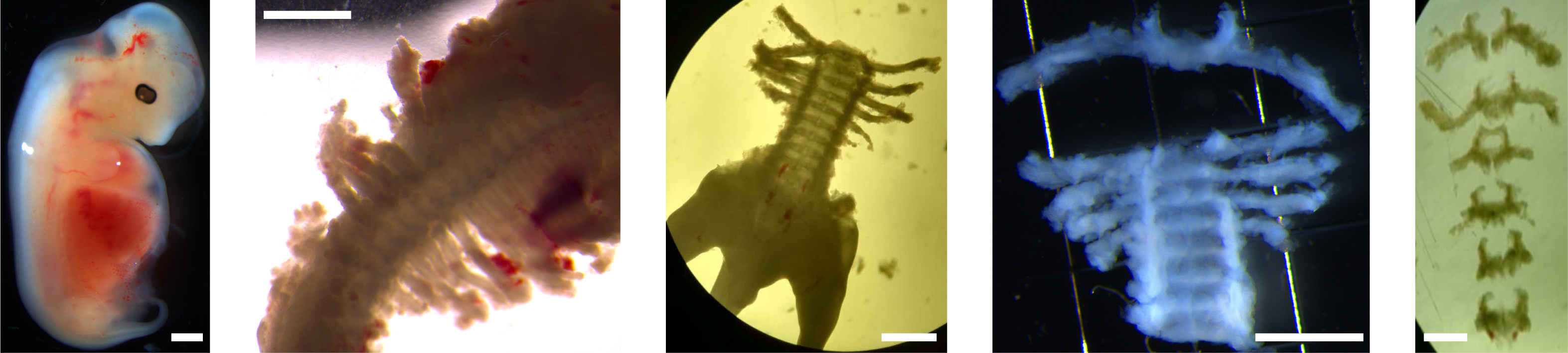

### Supplemental Figure 2

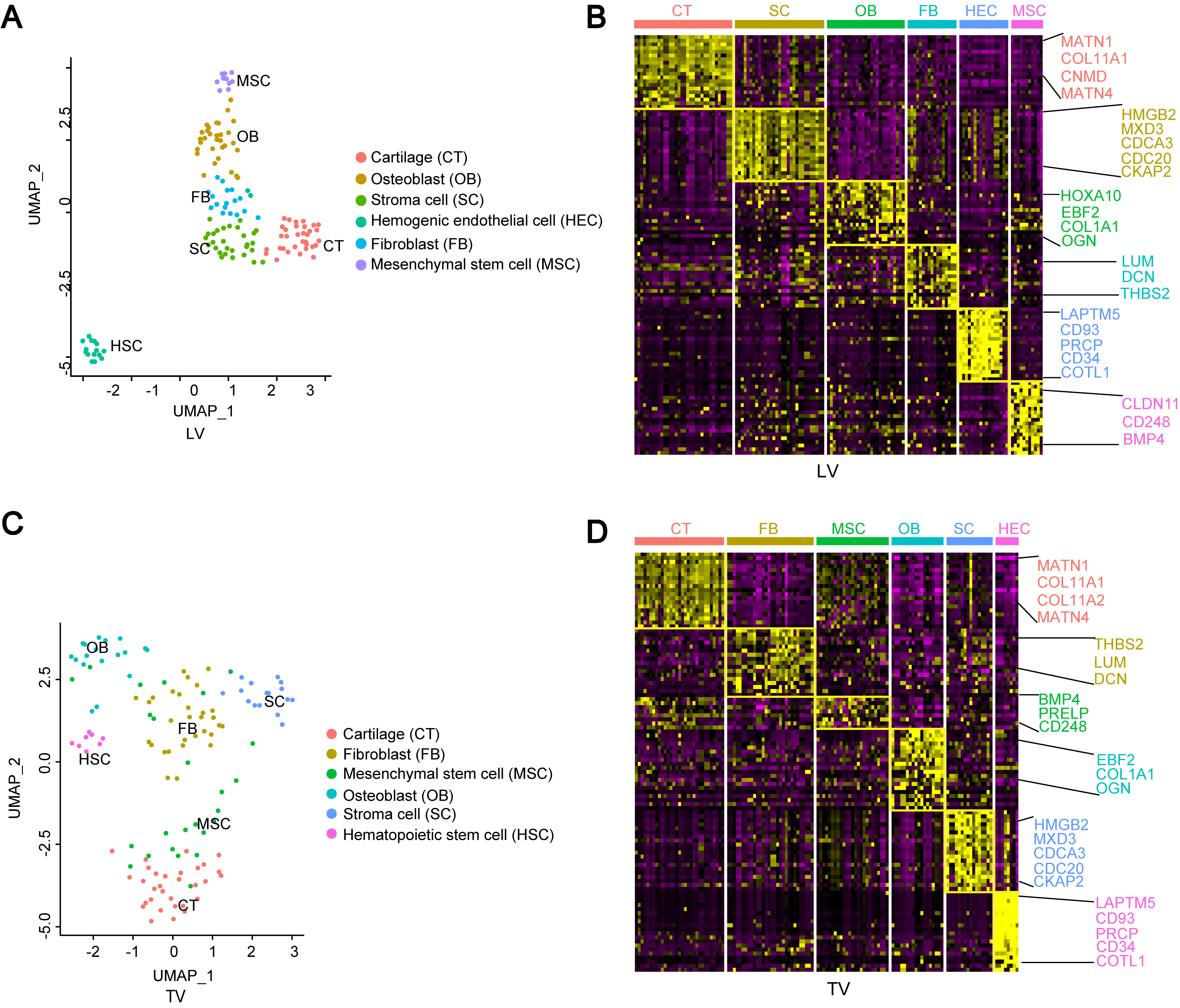
